# *Pontiella agarivorans* sp. nov., a novel marine anaerobic bacterium capable of degrading macroalgal polysaccharides and fixing nitrogen

**DOI:** 10.1101/2023.06.09.544357

**Authors:** Na Liu, Veronika Kivenson, Xuefeng Peng, Zhisong Cui, Thomas S. Lankiewicz, Kelsey M. Gosselin, Michelle A. O’Malley, David L. Valentine

## Abstract

Marine macroalgae produce abundant and diverse polysaccharides which contribute substantially to the organic matter exported to the deep ocean. Microbial degradation of these polysaccharides plays an important role in the turnover of macroalgal biomass. Various members of the *Planctomycetes-Verrucomicrobia-Chlamydia* (PVC) superphylum are degraders of recalcitrant polysaccharides in widespread anoxic environments. In this study, we isolated a novel anaerobic bacterial strain NLcol2^T^ from microbial mats on the surface of marine sediments offshore Santa Barbara, California, USA. Based on 16S rRNA gene and phylogenomic analyses, strain NLcol2^T^ represents a novel species within the *Pontiella* genus in the *Kiritimaitellaeota* phylum (within the PVC superphylum). Strain NLcol2^T^ is able to utilize various monosaccharides, disaccharides, and macroalgal polysaccharides such as agar and iota-carrageenan. A near-complete genome also revealed an extensive metabolic capacity for anaerobic degradation of sulfated polysaccharides, as evidenced by 202 carbohydrate-active enzymes (CAZymes) and 165 sulfatases. Additionally, its ability of nitrogen fixation was confirmed by nitrogenase activity detected during growth on nitrogen-free medium, and the presence of nitrogenases (*nifDKH*) encoded in the genome. Based on the physiological and genomic analyses, this strain represents a group of bacteria that may play an important role in the degradation of macroalgal polysaccharides and with relevance to the biogeochemical cycling of carbon, sulfur, and nitrogen in marine environments. Strain NLcol2^T^ (= DSM 113125^T^ = MCCC 1K08672) is proposed to be the type strain of a novel species in *Pontiella* genus, and the name *Pontiella agarivorans* sp. nov. is proposed.

**Importance:** The growth and preservation of marine macroalgae is considered as a carbon dioxide reduction strategy. However, we still have limited knowledge about the eventual fate of carbon stored in macroalgae. Diverse heterotrophic microbial communities in the ocean specialize on these complex polymers, for example, members in the *Kiritimatiellaeota* phylum. However, only four type strains within the phylum have been cultured and characterized to date, and there is limited knowledge about the metabolic capabilities and functional role of this phylum in the environment. The new isolate strain NLcol2^T^ expands the known substrate range of this phylum with the capability to utilize macroalgal polysaccharides agar and iota-carrageenan. It is also the first strain in the *Kiritimatiellaeota* phylum to be reported with nitrogen fixing ability.

## 1 Introduction

Marine macroalgae are important primary producers in coastal ecosystems. They sequester about 173 TgC yr^-1^ into their biomass and are considered as part of the “blue carbon” in the ocean (1). Seaweed cultivation has been considered as one of the promising strategies to mitigate the increasing amount of anthropogenic CO_2_ and climate change (National Academies of Sciences, Engineering, and Medicine, 2022). A recent study shows that 24% of macroalgae will eventually reach the seafloor and thus export the fixed carbon to the deep ocean (3). However, there are still uncertainties as to the fate of the blue carbon fixed into macroalgal biomass after sinking to the seafloor. Although many marine animals can digest macroalgae, the polymers within macroalgal biomass will eventually be degraded by heterotrophic microbes specialized on degrading recalcitrant organic matter.

Polysaccharides are important structural and storage compounds in macroalgae. They are highly diverse due to various monosaccharide units, glycosidic linkages, as well as different moieties (e.g., sulfate, amino, methyl, or acetyl groups) added to the carbohydrate backbone (4). In contrast to terrestrial plants, marine macroalgae produce polysaccharides decorated by sulfate and other functional groups, for example, agars, porphyrans, and carrageenans extracted from red algae, laminarin, fucoidans, and alginates from brown algae, ulvans from green algae, etc. (5–7). Agar is a mixture of agarose and agaropectin which is commonly used as solidifying agent for culture media. Agarose is composed of alternating α-1,3 linked D-galactose and β-1,4 linked 3,6-anhydro-α-L-galactose with little sulfate modification, while agaropectin is heavily modified with sulfate (8–10). Carrageenan is structurally related to agarose, except the β-linked unit is D-galactose-6-sulfate (9). Fucoidan is also a sulfated polysaccharide composed mainly of L-fucose units adorned with sulfate esters, while minor xylose, galactose, mannose, glucuronic acid can be present too (11).

Microbial degradation of macroalgal polysaccharides involves complex metabolic pathways and requires a large number of enzymes during the process (12–15). For example, hundreds of enzymes were shown to be involved in fucoidan degradation in *Verrucomicrobia* (14). Classic genomic features for glycan degradation were known as polysaccharide utilization loci (PULs) found in *Bacteroidetes* bacteria, especially the starch utilization system (Sus) of *Bacteroides thetaiotaomicron* (16, 17). They contain co-regulated gene clusters of cell surface glycan-binding lipoproteins for the detection of carbohydrates (typically *sus*D homologs), TonB-dependent transporters (typically *sus*C homologs) for the uptake of molecules into the cell, carbohydrate-active enzymes (CAZymes) for substrate-specific degradation, as well as sensors and transcriptional regulators. CAZymes, especially glycoside hydrolases (GHs) and polysaccharide lyases, can break down polysaccharides into oligosaccharides (18). Algal polysaccharide degradation has been well studied in *Zobellia galactanivorans* Dsij^T^, the marine *Bacteroidetes* model for the discovery of agarases, porphyranases, and carrageenases (12, 13, 19). As most marine polysaccharides are sulfated, another group of enzymes called sulfatases are needed in the degradation pathway, which can cleave sulfate ester groups off the carbohydrate backbone (20). Sulfatases are activated via post-translational modification by other enzymes before functioning. The most common one is formylglycine-generating enzyme (FGE) which transforms a cysteine or serine residue into a catalytic formylglycine (21). These fGly-sulfatases are classified as type I sulfatases (family S1) which contain all carbohydrate sulfatases and is the largest sulfatase family (7). Although less well studied, the anaerobic sulfatase-maturing enzyme can mature either cysteine or serine sulfatases under anaerobic conditions (22).

Members of the PVC superphylum (named for *Plantomycetes, Verrucomicrobia, Chlamydiae)* include degraders of recalcitrant glycopolymers, though much of their true functional diversity has been obscured by the lack of cultivated representatives (23–26). The PVC superphylum also consists of phyla *Kiritimatiellaeota* and *Lentisphaerae* as well as uncultured candidate phyla from environmental samples (27). Though highly diverse biologically and ecologically, they together form a deeply rooted monophyletic group on the basis of 16S rRNA gene analysis (28). The *Kiritimatiellaeota* phylum was established in 2016 and was previously recognized as the Subdivision 5 of *Verrucomicrobia* in the PVC superphylum (29). The geographic distribution of 16S rRNA gene sequences reveals that bacteria in phylum *Kiritimatiellaeota* are common to anoxic environments ranging from the intestine of animals to hypersaline sediments and wastewater (29). However, there are only four cultivated strains reported to date, and we know little about their metabolic capabilities and functional role in the environment. The first cultivated strain, *Kiritimatiella glycovorans* L21-Fru-AB^T^, is a halophilic saccharolytic bacterium isolated from an anoxic cyanobacterial mat from a hypersaline lake on the Kiritimati Atoll (30). *Pontiella desulfatans* F1^T^ and *Pontiella sulfatireligans* F21^T^ were isolated from Black Sea sediments and are capable of degrading sulfated polysaccharides like iota-carrageenan and fucoidan (31, 32). Unlike these three obligate anaerobes, *Tichowtungia aerotolerans* S-5007^T^ was isolated from surface marine sediment and can grow under microaerobic conditions (33).

Macroalgal polysaccharides are carbon rich, but depleted in nutrients including nitrogen and phosphorus. Among other isolates of *Kiritimatiellaeota* ammonium has been identified as the nitrogen source, but nitrogen fixation has not been observed. Mo-dependent nitrogenase is the most common and widely studied enzyme that perform nitrogen fixation. It contains two components: an Fe protein as the reductase (*nifH*) collecting and transferring electrons, and a MoFe protein (*nifDK*) which binds dinitrogen (N_2_) and converts it to ammonia (NH_3_) (34). Nitrogenases are highly oxygen-sensitive, but even though there are diverse anaerobes in the PVC superphylum, only a few studies demonstrated nitrogen fixation in this superphylum (35–38) and no reports in the *Kiritimatiellaeota* phylum. Moreover, we have little knowledge as to where *nif* genes were acquired from by the nitrogen-fixing members in the PVC superphylum.

In this study, we enriched and isolated a novel anaerobic bacterial strain NLcol2^T^ in the *Kiritimatiellaeota* phylum with agar as substrate and nitrogen gas as the sole nitrogen source. We characterized this strain by phylogenomic, morphological, chemotaxonomic, and physiological traits. We further investigated its metabolic potential by analyzing CAZymes, sulfatases and nitrogenases in the genome in detail.

## 2 Materials and Methods

### 2.1 Inoculum source, enrichment, and isolation of strain NLcol2^T^

Strain NLcol2^T^ was enriched and isolated from microbial mats found on the surface of marine sediments at Shane Seep (34.40616 N, 119.89047 W) within the Coal Oil Point seep field offshore Santa Barbara, California, USA. Microbial mat samples were collected at 20 m depth with an in-situ temperature of 15 °C in October 2017. The seep area is characterized by a large amount of hydrocarbon gas emissions, microbial mat coverage, and high sulfide and alkalinity in sediment porewater (39–41). The samples used for inoculum contained both microbial mats and partially decomposed macroalgae (**Figure S1a**). The microbial mats were scraped off their attached surface as the inoculum source. The cultures were enriched anaerobically in semi-solid agar (0.25% w/v) in the top layer of the sulfide gradient media (**Figure S1b**) modified from Kamp et al., 2006. Cultures were maintained at room temperature in the dark and were transferred into fresh media every two to three weeks for a year. A preliminary survey of microbial communities by 16S rRNA amplicon sequencing revealed a novel bacterium being dominant in the enrichment cultures.

Further isolation of strain NLcol2^T^ was performed by streaking on agar plates in an anaerobic chamber (Coy Laboratory Products) (**Figure S1c**). The medium is the same as the top agar medium in enrichment cultures, except that 1.5 % w/v agar was added as both gelling agent and substrate and 2 mM sulfide added as reducing agent. The Petri dishes were kept in the anaerobic chamber at room temperature (22 °C). Single colonies formed after three weeks and were picked from agar plates. Streak plating was repeated for three more rounds to ensure the purity of the culture. Pure culture was subsequently maintained in liquid media with D-galactose (1g/L) as substrate at 22 °C and was transferred every other week. A full modified medium contained: 28.0 g NaCl, 10.0 g MgCl_2_ ·6 H_2_O, 3.8 g MgSO_4_ ·7 H _2_O, 0.6 g CaCl_2_ ·2 H _2_O, 1.0 g KCl, 37 mg K_2_HPO_4_, 4 mg Na_2_MoO_4_, 50 mg Na_2_S_2_O_5_, 2 mg FeCl_3_ ·6 H _2_O, 10.0 mL modified Wolin’s Mineral Solution (see DSMZ medium 141), 0.5 mL Na-resazurin solution (0.1% w/v), 1.0 g D-galactose, 1.0 g NH_4_Cl (optional), 0.75 g Na_2_CO_3_, 0.5 g Na_2_S· 9 H_2_O, 10.0 mL Wolin’s Vitamin Solution (see DSMZ medium 141), in 1000 mL distilled water. All ingredients except carbonate, sulfide and vitamins were dissolved under N_2_/CO_2_ (80:20) atmosphere in Hungate tubes or serum bottles and autoclaved. Carbonate was added from a sterile anoxic stock solution prepared under N_2_/CO_2_ (80:20) atmosphere. Sulfide and vitamins were added from sterile anoxic stock solutions prepared under 100% N_2_ gas. Purity of the isolate was checked by full-length 16S rRNA gene sequencing and observation of morphology under the microscope.

### 2.2 Phylogenetic reconstruction by 16S rRNA gene

Full-length 16S rRNA gene of strain NLcol2^T^ was sequenced by GENEWIZ (Azenta Life Sciences), from colonies grown on agar plates. 16S rRNA gene sequence was searched using the website tool BLASTn (42) against the 16S rRNA database and compared to the sequence identity to the other four isolated strains in the *Kiritimatiellaeota* phylum.

To construct a phylogenetic tree based on the 16S rRNA gene, 106 sequences over 1200 bp from the *Kiritimatiellales* order in SILVA Ref NR SSU r138.1 database (released August 2020, accessed November 2021) (43) were selected for alignment. The full-length 16S rRNA genes of strain NLcol2^T^, *Tichowtungia aerotolerans* strain S-5007^T^, and two *Verrucomicrobia* (ABEA03000104, AF075271 as outgroups) were also added to the alignment using SINA Aligner v1.2.11 (44). The alignment was trimmed using the “gappyout” method in TrimAl v1.4 (45) to remove ambiguous ends and columns with >95% gaps. All trimmed nucleotide sequences represent >50% of the 1568 alignment columns. A maximum-likelihood tree was constructed using RAxML v.8.2.9 (46) with GTRGAMMA model of evolution. Rapid bootstrap search was stopped after 1000 replicates with MRE-based criterion. The best-scoring ML tree with support values was visualized in the iTOL server (47).

### 2.3 Genome sequencing and analyses

Genomic DNA was extracted from the isolate cultures using FastDNA Spin Kit for Soil (MP Biomedicals, OH). Genomic DNA library preparation and sequencing were performed at the University of California Davis Genome Center on Illumina HiSeq 4000 platform with 150-base pair (bp) paired-end reads. Trimmomatic v.0.36 (48) and Sickle v.1.33 (49) were used to remove adapter and low quality or short reads. Trimmed reads were assembled into contigs using MEGAHIT v.1.1.1 (50). Contigs longer than 2500 bp were kept and the trimmed reads were mapped back to those contigs using Bowtie2 v.2.3.4.1 (51) and Samtools v.1.7 (52). Contigs were visualized using Anvi’o v.3 interactive interface (53) and manual binning was performed based on coverage, GC content, tetranucleotide frequency signatures. Completion and redundancy for the reconstructed genome was determined using CheckM v.1.0.7 (54).

Open reading frame (ORF) features and protein-coding gene sequences were predicted using Prodigal v.2.6.3 (55). Annotation was assigned to proteins using hmmer v.3.1b2 (56) hmmscan searching against the Pfam v.32.0 (57) and TIGRFAMs v.15.0 (58) databases with a maximum e-value of 1×10^-7^. Information on protein family, domain and conserved site were confirmed using InterProScan5 (59). The amino acid sequences of protein-coding genes were further searched against NCBI’s Conserved Domain Database (CDD) (60) using the RPS-BLAST program v.2.7.1. The cdd2cog script (61) was used to assign COG (Cluster of Orthologous Groups) categories (62) to each protein-coding gene. Protein sequences were also submitted to the BlastKOALA server (63) for KEGG Orthology (KO) ID assignments. Ribosomal RNA genes were determined by RNAmmer v.1.2 (64). tRNA genes were predicted by tRNAscan-SE 2.0 server (65). Metabolic pathways were reconstructed using KEGG Mapper (66) and MetaCyc database (67).

For phylogenomic analyses, 24 high-quality genomes in the *Kiritimatiellales* order from NCBI’s GenBank database and the Genome Taxonomy Database (GTDB) r95 were selected (accessed on Feb 1, 2021). *Opitutus terrae* PB90-1 from the *Verrucomicrobia* phylum was selected as the outgroup. All genomes meet the GTDB quality criterion based on completeness and redundancy from CheckM: completeness – 5×redundancy > 50. 120 single-copy genes were searched and aligned using GTDB-Tk v1.4.0 (68). The concatenated alignment was further trimmed using TrimAl v1.4 (45) with “gappyout” parameter, which results in a final alignment with 4488 amino acid columns. Maximum-likelihood phylogenetic tree was calculated using RAxML v.8.2.9 (46) with PROTGAMMALG model of evolution. Rapid bootstrap search was stopped after 350 replicates with MRE-based criterion. The best-scoring ML tree with support values was visualized in the iTOL server (47). The average nucleotide identity (ANI) and average amino acid identity (AAI) between genomes were calculated using the ANI/AAI calculator (69).

Carbohydrate-active enzymes were predicted using dbCAN2 meta server (70). In brief, uploaded protein sequences were searched against the dbCAN CAZyme domain HMM database v.7, CAZy database (www.cazy.org) and PPR library using HMMER, DIAMOND, and Hotpep programs respectively (70). Only genes predicted by no less than two programs were defined as CAZymes for further analysis. CAZyme gene clusters were predicted by the CGC-Finder on dbCAN2 server with at least one CAZyme and one transporter detected within a maximum distance of two genes (70). To classify sulfatases into families and subfamilies, gene sequences with an annotated sulfatase domain (PF00884) were searched and classified by the SulfAtlas database v.1.1 (20) using the BLASTp program (42). Additionally, SignalP v.5 (71) was used to predict signal peptides for translocation of sulfatases into the periplasmic space and outside of the cells.

To better understand the evolution of nitrogen fixation in the *Kiritimatiellaeota* phylum, reannotation and phylogenetic analysis of the *nifH* gene were performed for all 52 genomes in this phylum from NCBI’s GenBank database (accessed on Mar 3, 2020). The same annotation pipeline described above was used to keep consistency and allow better comparison. *nifH* gene sequences were aligned with 879 full-length *nifH* genes from the genomes of cultivated diazotrophs (https://www.zehr.pmc.ucsc.edu/Genome879/) using MUSCLE v.3.8 (72). Two light-independent protochlorophyllide reductases were included as outgroups: ChlL from *Trichormus variabilis* ATCC 29413 (WP_011320185.1) and BchL from *Chlorobium limicola* DSM 245 (WP_012467085.1). The alignment was trimmed in Jalview v.2.10.5 (73) to remove ambiguous ends and the columns with >95% gaps. All trimmed amino acid sequences represent >81% of the alignment columns. A maximum-likelihood tree was constructed using RAxML v.8.2.9 (46) with LG substitution model plus GAMMA model of rate heterogeneity. Rapid bootstrap search was stopped after 350 replicates with MRE-based criterion. The best-scoring ML tree with support values was visualized in the iTOL server (47).

### 2.4 Microscopy

Bacterial cells were observed under the Olympus BX51 Upright Compound Microscope at the Neuroscience Research Institute, University of California Santa Barbara. Oil and differential interference contrast (DIC) microscopy were applied for better resolution and contrast.

To obtain high-resolution images, cell morphology was examined under the transmission electron microscope (TEM). For TEM imaging, cells grown on the agar plates were fixed with modified Karnovsky’s fixative (2% paraformaldehyde and 2.5% glutaraldehyde in 0.1 M sodium phosphate buffer) and spun down into a cell pellet. Cells were rinsed in 0.1 M sodium phosphate buffer and fixed again with 1% osmium tetroxide in the same buffer. After another rinse, they were dehydrated in 50% EtOH, 75% EtOH, 95% EtOH, 100% EtOH and propylene oxide twice. Cells were pre-infiltrated in 1:1 propylene oxide:resin (Epon/Alradite mixture) overnight, infiltrated in 100% resin and embedded in fresh resin at 60 °C overnight. Ultrathin sections were cut using a Diatome diamond knife. Sections were picked up on copper grids and imaged in a FEI Talos 120C transmission electron microscope at the Biological Electron Microscopy Facility, University of California Davis.

### 2.5 Growth tests

Bacterial growth of strain NLcol2^T^ was monitored by measuring optical density (OD) of liquid cultures at 600 nm wavelength. Growth at different temperature (4, 10, 14, 22, 26, 31, 37, 55 °C), salinity (0%, 1%, 2%, 2.5%, 3%, 4%, 5%, 6% NaCl) and pH (4.0, 5.0, 5.5, 6.0, 6.5, 7.0, 8.0, 9.0) conditions were determined in triplicates when growing on D-galactose with ammonium supplied. Growth was tested on various substrates (1 g/L) in triplicates at optimum temperature, salinity and pH conditions with ammonium supplied: D-glucose, D-galactose, D-fructose, L-fucose, L-rhamnose, D-mannose, D-mannitol, meso-inositol, D-arabinose, D-xylose, D-cellobiose, lactose, sucrose, maltose, xylan, starch, cellulose, alginic acid, agarose, agar, iota-carrageenan, fucoidan from *Macrocystis pyrifera*, dried red algae (*Porphyra* spp.), dried brown algae (*Saccharina japonica* and *Saccharina latissima*), and the giant kelp (*Macrocystis pyrifera*).

### 2.6 Metabolite analysis

To quantify the metabolic products of strain NLcol2^T^ from galactose fermentation, cultures were grown in triplicates at room temperature (22 °C) with D-galactose as carbon source for 10 days. No ammonium was added in the media and N_2_ gas was served as the sole nitrogen source. Growth was monitored by measuring optical density (OD) at 600 nm wavelength. 2mL of culture was subsampled each day (twice a day during exponential phase) for quantification of metabolites.

The chromatography protocols used in this study are similar to those previously described for Gilmore et al., 2019 and Peng et al., 2021. Galactose, acetate, succinate, fumarate and malate concentrations were measured on an Agilent Infinity 1260 (Agilent Technologies, Santa Clara, CA, USA) high-performance liquid chromatograph (HPLC) using an Aminex HPX-87H analytical column (part no. 1250140, Bio-Rad, Hercules, CA, USA) protected by, first, a 0.22 µm physical filter, followed by a Coregel USP L-17 guard cartridge (Concise Separations, San Jose, CA, USA). Separations were performed at 60 °C with a flow rate of 0.6 mL/min and a 5 mM sulfuric acid (H_2_SO_4_) mobile phase. Acetate, succinate, fumarate, and malate were measured using a variable wavelength detector set to 210 nm, while galactose was measured using a refractive index detector set to 35 °C. Samples and standards for HPLC were acidified to a concentration of 5 mM H_2_SO_4_, incubated for 5 min at room temperature, and spun at maximum speed in a tabletop centrifuge for 5 min to pellet bacterial cells. The samples were removed from above the cell pellet, and 0.22 µm filtered through a polyethersulfone (PES) membrane into HPLC vials with 300 µL polypropylene inserts. Standard curves for each compound of interest were constructed using triplicate standards of 0.1, 0.5, and 1.0 g/L. Peaks were integrated using OpenLab CDS analysis software (version 2.6, Agilent Technologies).

Hydrogen gas production was measured on a Fisher Scientific TRACE 1300 Gas Chromatograph (Thermo Fisher Scientific, Waltham, MA) using a TRACE TR-5 GC Column (part no. 260E113P, Thermo Fisher Scientific) at 30 °C, with an Instant Connect Pulsed Discharge Detector (PDD) (part no. 19070014, Thermo Fisher Scientific) at 150 °C, and ultra-high purity He as a carrier gas. All injections of samples and standards were 100 µL in volume. Supplier-mixed standards of 50 ppm, 500 ppm, and 1% hydrogen were run before and after injecting samples, and hydrogen peaks were integrated using Chromeleon Chromatography Data System (CDS) Software (version 6.8, Thermo Fisher Scientific). CO_2_ was not considered due to the carbonate-buffered medium and N_2_/CO_2_ atmosphere.

### 2.7 Cellular fatty acid analysis

The cellular fatty acid composition of strain NLcol2^T^ was determined from cells grown at 22 °C to late-log phase in liquid medium with 1.0 g/L D-galactose as carbon source and nitrogen gas as nitrogen source. Cells were centrifuged down at 10,000 × g for 10 mins and were frozen in -80 °C.

Cellular fatty acids were extracted twice using a modified Folch method (76) with a chloroform: methanol mixture (2:1) and tridecanoic acid as an internal standard. The samples were partitioned and the organic phase containing the total lipid extract (TLE) was retained. Transesterification of the TLE was performed by adding toluene and 1% sulfuric acid in methanol to the TLE after it was brought to complete dryness under N_2_. The acidic methanol/toluene TLE was heated at 90 °C for 90 minutes to produce fatty acid methyl esters (FAME). The FAMEs were extracted from the acidic methanol by adding hexane and water, vortexing, centrifuging, and removing the top (hexane) fraction to a new vial twice. The combined transesterified hexane extracts were dried under N_2_ to a final volume of 300 μL. Each extract was spiked with methyl heptadecanoate to calculate the recovery of the internal standard and analyzed by GC-FID.

Concentration analysis was done on a gas chromatograph flame ionization detector (GC-FID), HP 5890 Series II GC-FID. Chromatography was performed with a 30 m × 0.25 mm internal diameter (ID), 0.25 μm pore size, fused silica Omegawax capillary column with a splitless 1-μL injection. Initial oven temperature was set at 50 °C and held for 2 min, followed by a 10 °C min ^−1^ ramp to 150 °C, then a 5 °C min ^−1^ ramp to the final temperature of 265 °C. A certified reference material (FAME 37, Supelco) was run to calculate retention times and identify peaks. Peak identification was further confirmed from their mass spectra.

### 2.8 Acetylene reduction assay

To test the nitrogenase activity of strain NLcol2^T^ when growing with nitrogen gas as the sole nitrogen source, acetylene reduction assay was performed following Hardy et al., 1968. In short, acetylene (C_2_H_2_) can be reduced to ethylene (C_2_H_4_) when nitrogenases actively fix nitrogen gas at the same time. Cultures were grown on D-galactose in triplicates at 22 °C and triplicate media bottles without inoculation were used as controls. 1.2 mL of acetylene was injected to all culture and control bottles, which contained 80 mL of liquid and 80 mL of headspace pressurized at 150 kPa at the beginning. Gas concentrations and OD_600_ were measured at 6 time points during the 18-day incubation. Acetylene and ethylene concentrations were resolved on a Shimadzu 8A Gas Chromatograph with a flame ionization detector (GC-FID). 1.5 mL samples and standards were injected, then carried by N_2_ at a flow rate of 20 mL/min through an n-octane on Res-Sil C packed column (Restek, Centre County, PA, USA) set at 25 °C. 0.5% and 1.0% GASCO calibration gas mixtures of acetylene and ethylene (Cal Gas Direct Incorporated, Huntington Beach, CA, USA) were used for the standard curves.

## 3 Results and Discussion

### 3.1 Phylogenetic analyses

Phylogenetic placement of strain NLcol2^T^ was determined by comparing full-length 16S rRNA gene, single-copy genes, and whole-genome similarity metrics including average nucleotide identity (ANI) and average amino acid identity (AAI).

Strain NLcol2^T^ was classified within the R76-B128 clade of the *Verrucomicrobia* phylum under the current SILVA taxonomy (SILVA Ref NR SSU r138.1). Full-length 16S rRNA gene of the isolate shares 84.1%, 88.9%, 92.9%, and 94.5% identity with the four reported cultivated strains in the proposed *Kiritimatiellaeota* phylum: *Kiritimatiella glycovorans* strain L21-Fru-AB^T^, *Tichowtungia aerotolerans* strain S-5007^T^, *Pontiella sulfatireligans* strain F21^T^, and *Pontiella desulfatans* strain F1^T^, respectively. Strain NLcol2^T^ is more closely related with *Pontiella* strains F1 and F21 than *K. glycovorans* and *T. aerotolerans*. The 16S rRNA gene identities compared to *Pontiella* strains F1 and F21 were absolutely higher than the 86.5% threshold for family level, but fall on the edge of the threshold for a new genus as 94.5% (77). A maximum-likelihood tree of 16S rRNA gene sequences from the *Kiritimatiellaeota* phylum was reconstructed by RAxML (**Figure S2**). The R76-B128 clade formed a monophyletic group with MSBL3 clade as the sister group, both of which are in a different cluster from the *Kiritimatiellaceae* family. It is clear that strain NLcol2^T^ is not affiliated with *K. glycovorans* within the *Kiritimatiellaceae* family nor with *T. aerotolerans* within the MSBL3 clade, but belongs to the R76-B128 clade within the *Kiritimatiellales* order as *Pontiella* strains F1 and F21 (31). However, the bootstrap values in the R76-B128 clade were low and we cannot determine whether strain NLcol2^T^ represents a novel genus or belongs to the same *Pontiella* genus as strains F1 and F21 by 16S rRNA gene phylogeny alone.

To resolve the phylogeny of strain NLcol2^T^ in detail, we further performed genome-level phylogenetic analyses using the Genome Taxonomy DataBase toolkit (GTDB-Tk). A concatenated phylogenomic tree was reconstructed from 120 bacterial single-copy genes of 25 genomes in the *Kiritimatiellales* order (**Figure 1**). Here, strain NLcol2^T^ falls within the proposed *Pontiella* genus with a bootstrap value of 100. Additionally, the average amino acid identity (AAI) values within the genomes of the proposed *Pontiella* genus is 68.43% (**Table S1**), which is slightly above the threshold of 65% for same genus (78). However, within *Pontiella* genus, it represents a different group from the one including strains F1 and F21. The average nucleotide identity (ANI) values of the genomes between strain NLcol2^T^ and strains F21 and F1 are 72.73% and 73.71% respectively, which was much lower than the 95% ANI criterion for the same species (79, 80). Therefore, we propose that strain NLcol2^T^ represents a novel species within the *Pontiella* genus according to the phylogenetic analyses above.

**Figure 1.**
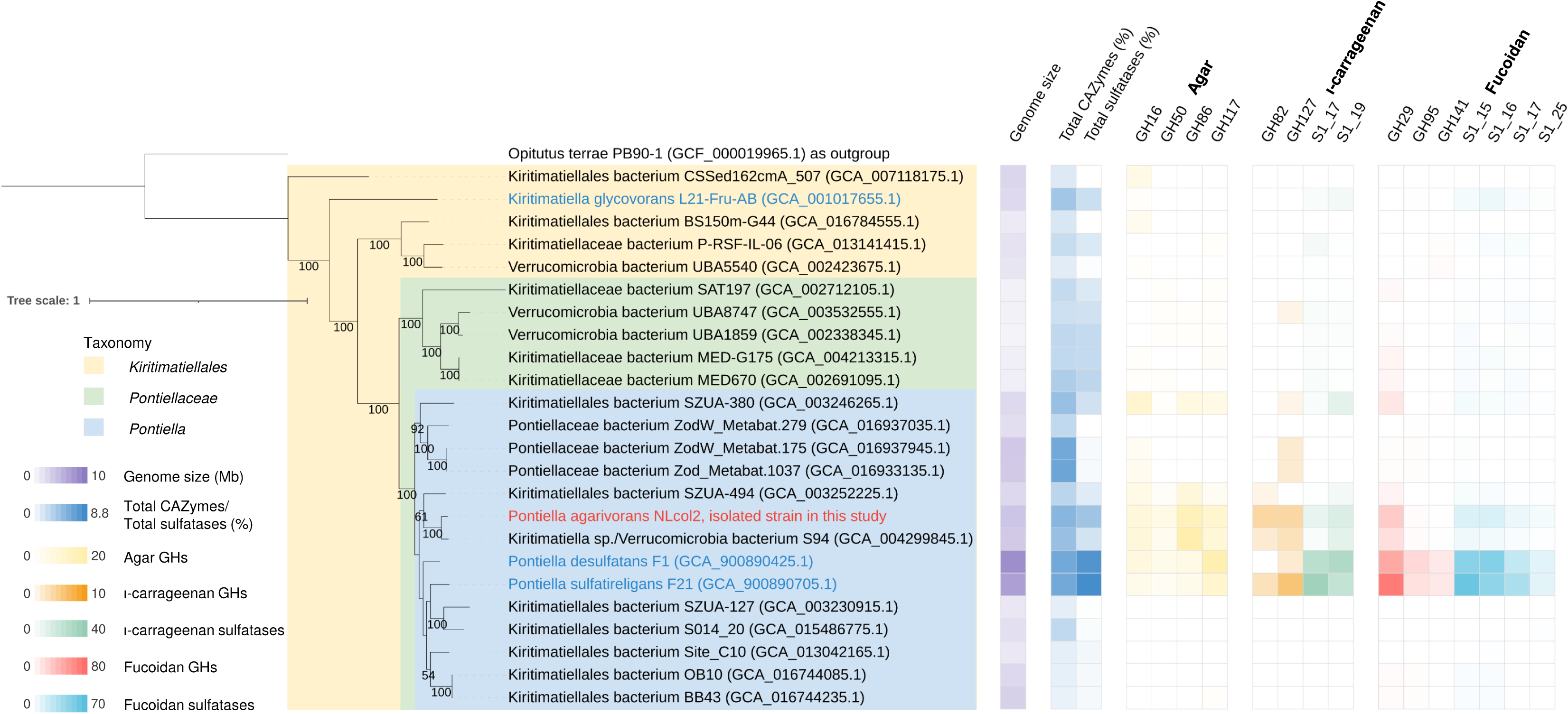
Concatenated maximum-likelihood phylogenomic tree of 120 bacterial single-copy genes from 24 genomes in the *Kiritimatiellales* order. Strain NLcol2 is labeled in red and other cultivated strains are labeled in blue. A *Verrucomicrobia* genome was selected as outgroup and the tree was rooted there. Bootstrap values over 50 are shown on the nodes. Genome size (Mb), total CAZymes (%) and total sulfatases (%) as a percentage of all protein-coding genes in each genome are presented as reference. The number of glycoside hydrolases (GH) and sulfatase homologs involved in the degradation pathways of agar, iota-carrageenan, and fucoidan are presented in the heatmap.

### 3.2 General features of the genome

The draft genome of strain NLcol2^T^ is 95% complete with 4% redundancy. The genome consists of 12 contigs (N50 is 1,265,434 bp) with a total length of 4,436,865 bp and the mean coverage is 593x. DNA G+C content is 52.4 mol%. 5S, 16S and 23S rRNA genes and 50 tRNA genes were found in the genome.

3,611 ORFs were predicted by Prodigal, among which 2,757 proteins in the genome were assigned with COG functional category codes. The number of genes in each functional category is shown in **Figure S3**. The greatest number of genes (357) fall into the category of inorganic ion transport and metabolism. More genes are involved in carbohydrate (260) and amino acid (188) transport and metabolism than those of nucleotides (65) and lipids (61), which is similar to that in *Kiritimatiella glycovorans* (Spring et al., 2016). A further detailed analysis of genes involved in macroalgal polysaccharide degradation and nitrogen fixation is presented in sections 3.4 and 3.5.

### 3.3 Morphologic and chemotaxonomic characterization of strain NLcol2^T^

Single colonies on agar plates were white or ivory, circular, and smooth after growing anaerobically for 2 weeks at 22 °C. Bacterial cells of strain NLcol2^T^ have a round to ovoid shape with a size of 1 µm in diameter observed under microscope (**Figure 2a**). Cells divide by binary fission and genes of bacterial cell division complex including FtsZ family were present. No motility or flagella were observed, although a full set of genes coding for flagellar assembly was present in the genome. No spore formation was observed. A Gram-negative cell wall structure of outer membrane, periplasmic space and cytoplasmic membrane was shown by electron microscopy (**Figure 2b**). There are also genes coding for proteins involved in lipopolysaccharide export and peptidoglycan synthesis in the genome. Some bacteria in the PVC superphylum exhibit compartments inside the cells (81), but like other strains in the *Kiritimatiellaeota* phylum, no compartmentalization of the cytoplasm was observed in strain NLcol2^T^. There were unknown inclusions or granules present inside the cells, and genes involved in the synthesis and utilization of polyphosphate and glycogen were found in the genome, which may serve as phosphate and energy storage materials, respectively.

**Figure 2.**
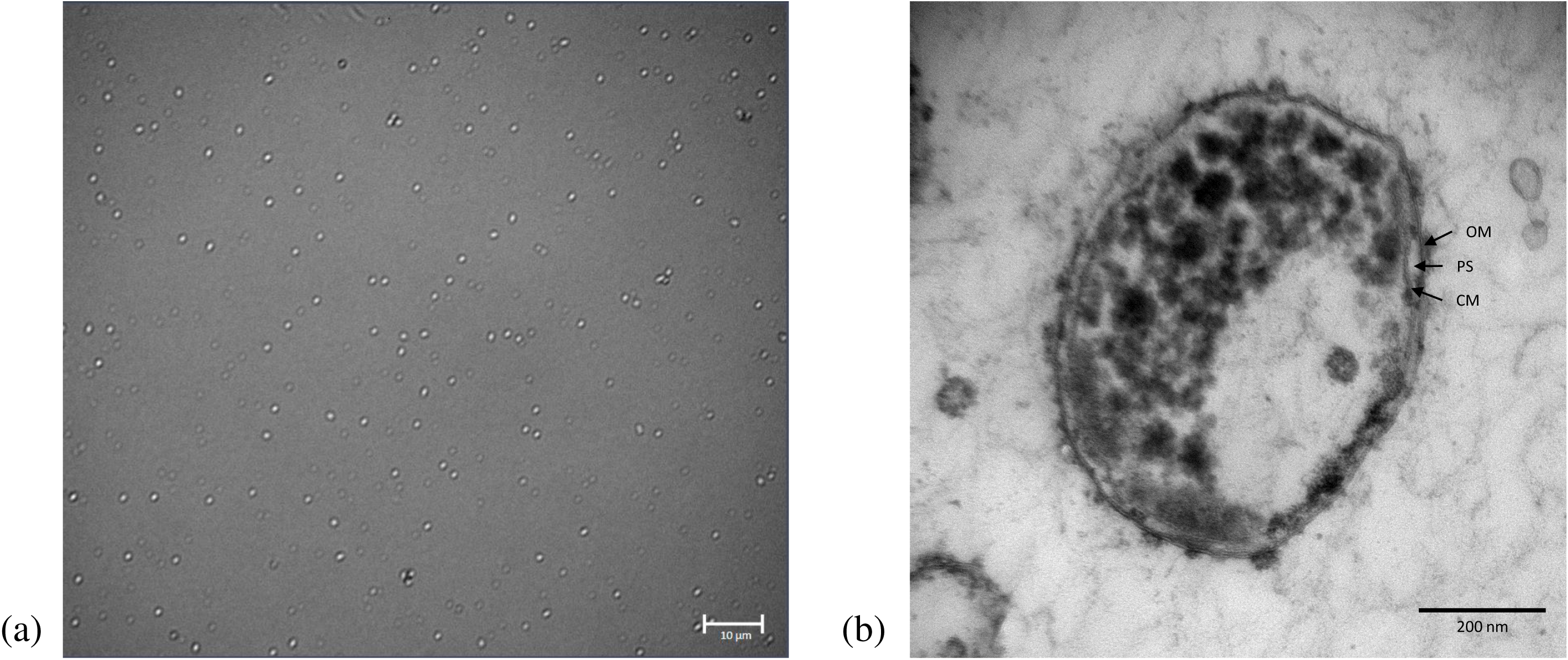
Microscopic images of strain NLcol2. (a) Bright field microscopy image; (b) Transmission electron microscopy image. OM, outer membrane; PS, periplasmic space; CM, cell membrane.

Major cellular fatty acids (>10% of total) of strain NLcol2^T^ include C18:0, *i-*C12:0, *i-*C18:0 and *i*-C14:0, in order of abundance. The major cellular fatty acid profile is quite different from *K. glycovorans* and *T. aerotolerans*, but almost the same as that in *Pontiella* strains F1 and F21, except that strain F21 also has *i*-C16:0 as one of the major components (**Table 1**). Again, this agrees with the phylogenetic placement of strain NLcol2^T^ in the *Pontiella* genus, being more closely related with strain F1 than strain F21. However, strain NLcol2^T^ can be further distinguished by a relatively higher composition of *i*-C18:0 than *i*-C14:0, while strain F1 has more *i*-C14:0 than *i*-C18:0 (**Table S2**). Other cellular fatty acids detected in strain NLcol2^T^ include C16:0, *i-*C16:0, C20:0, and *i-*C20:0 (**Table S2**).

**Table 1.** Comparison of phenotypic characteristics between strain NLcol2^T^ and four other isolated strains in the *Kiritimatiellaeota* phylum. Notations: NA, data not available. +, positive; -, negative; +/-, unstable, ceasing growth upon the second transfer. Data for strains other than NLcol2^T^ were referenced from literatures: a) van Vliet et al., 2020 (32); b) Spring et al., 2016 (29); c) Mu et al., 2020 (33). * Substrates were D-galactose for strain NLcol2^T^ and D-glucose for other strains. ** For strain S-5007^T^, acetate production was predicted from genomic data.

| Strains | *P. agarivorans*<br>NLcol2<sup>T</sup> | *P. desulfatans*<br>F1<sup>T a)</sup> | *P. sulfatireligans*<br>F21<sup>T a)</sup> | *K. glycovorans*<br>L21-Fru-AB<sup>T b)</sup> | *T. aerotolerans*<br>S-5007<sup>T c)</sup> |
| --- | --- | --- | --- | --- | --- |
| Isolation source | Microbial mat on marine sediment | Anoxic marine sediment | Anoxic marine sediment | Hypersaline microbial mat | Marine sediment |
| Cell shape | Spherical | Spherical | Spherical | Spherical | Spherical |
| Cell size (µm) | 1.0 | 0.5-1.2 | 0.5-1.0 | 1.0-2.0 | 0.5-0.8 |
| Motility | - | - | - | - | - |
| Genome size (Mbp) | 4.44 | 8.66 | 7.40 | 2.95 | 3.88 |
| DNA G+C content (mol%) | 52.4 | 56.0 | 54.6 | 63.3 | 53.1 |
| Major cellular fatty acids (>10% of total) | C18:0, *i*-C12:0, *i*-C18:0 | C18:0, *i*-C12:0, *i*-C14:0 | C18:0, *i*-C12:0, *i*-C18:0 | *i*-C14:0, C18:0 | C18:0, *i*-C12:0, *i*-C18:0, *i*-C16:0 |
| Growth Temperature (°C) |  |  |  |  |  |
| Range | 10-37 | 10-30 | 0-25 | 20-40 | 15-45 |
| Optimum | 31 | 25 | 25 | 28 | 33-35 |
| Growth Salinity (g/L NaCl) |  |  |  |  |  |
| Range | 10-60 | 10-31 | 10-50 | 20-180 | 5-80 |
| Optimum | 25-30 | 23 | 23 | 60-70 | 30-40 |
| Growth pH |  |  |  |  |  |
| Range | 6.0-9.0 | 6.5-8.5 | 6.0-8.5 | 6.5-8.0 | 6.0-8.5 |
| Optimum | 8.0 | 7.5 | 7.5 | 7.5 | 7.0-7.5 |
| Substrate utilization |  |  |  |  |  |
| Glucose | + | + | + | + | + |
| Galactose | + | + | + | +/- | + |
| Fructose | + | + | + | - | + |
| Fucose | - | + | + | - | NA |
| Rhamnose | - | + | + | +/- | + |
| Mannose | + | - | + | + | - |
| Mannitol | + | - | + | - | - |
| Arabinose | - | + | - | - | + |
| Xylose | + | + | + | + | - |
| Lactose | + | + | + | - | NA |
| Cellobiose | + | + | + | - | + |
| Sucrose | + | + | + | - | - |
| Maltose | + | + | + | - | - |
| Fucoidan | + | + | + | +/- | NA |
| iota-Carrageenan | + | - | + | +/- | NA |
| Xylan | + | - | - | NA | NA |
| Agar | + | - | - | - | - |
| Major non-gaseous fermentation products * | Succinate, acetate, malate | Acetate, ethanol, lactate | Acetate, ethanol, lactate | Ethanol, acetate | Acid (maybe acetate **) |
| Nitrogen sources | N<sub>2</sub>, NH<sub>4</sub><sup>+</sup> | NH<sub>4</sub><sup>+</sup> | NH<sub>4</sub><sup>+</sup> | NH<sub>4</sub><sup>+</sup> | NH<sub>4</sub><sup>+</sup> |

### 3.4 Physiology of growth

Strain NLcol2^T^ exhibited consistent growth between 10-37 °C (optimum 31 °C), with NaCl concentration between 10-60 g/L (optimum 25-30 g/L), and with pH 6.0-9.0 (optimum pH 8.0) when D-galactose was utilized as the substrate. It was determined as a mesophilic and neutrophilic bacterium, which is similar to the other four isolated strains from the *Kiritimatiellaeota* phylum (**Table 1**). Growth with ammonium supplied in the medium was faster than when dependent on nitrogen fixation. The doubling times are 15 h and 65 h when growing with and without ammonium respectively, at room temperature (22 °C). Strain NLcol2^T^ was considered as obligately anaerobic, being unable to grow with the presence of oxygen and even in non-reduced medium lacking sulfide as the reducing agent.

Strain NLcol2^T^ was able to grow on various carbohydrate substrates under optimal conditions with ammonium supplied, which includes D-glucose, D-galactose, D-fructose, D-mannose, D-mannitol, D-xylose, D-cellobiose, lactose, sucrose, maltose, xylan, agarose, agar, and iota-carrageenan. No growth was observed when supplied with L-fucose, L-rhamnose, D-arabinose, meso-inositol, starch, cellulose, alginic acid, or with fucoidan from *Macrocystis pyrifera*.

When growing on D-galactose, major fermentation products formed were succinate and acetate, with small amounts of malate and hydrogen gas also detected during the incubation (**Figure 3).** Initially, the culture was supplied with 4.71 ± 0.12 mM D-galactose with only 0.43 ± 0.06 mM D-galactose remaining following the 10-day incubation period. Taking all fermentation products into consideration, the fractional electron recovery for galactose fermentation by strain NLcol2^T^ was about 75%. A tentative fermentation balance was formulated as below:

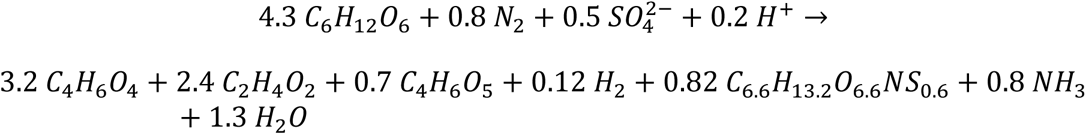

**Figure 3.**
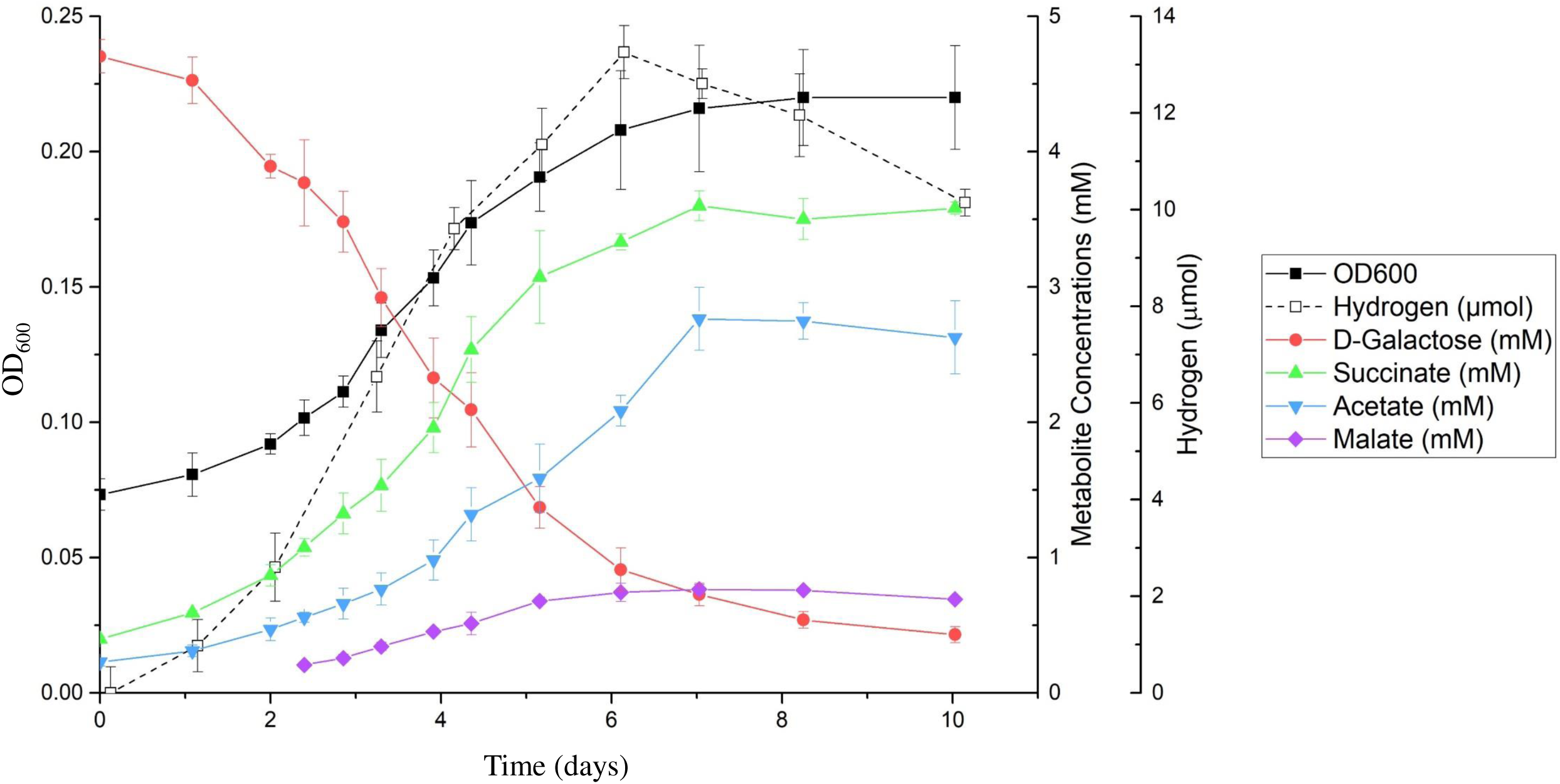
Time series of major metabolites from galactose fermentation by strain NLcol2. Growth curve was measured by optical density at 600nm (OD_600_).

### 3.5 Anaerobic degradation of macroalgal polysaccharides

#### 3.5.1 CAZyme analyses

202 genes (5.6% of all genes) were predicted to be CAZymes and associated carbohydrate-binding modules (CBM) by dbCAN2 meta server (70) (**Table S3**). Among these, 164 genes were annotated to be in the glycoside hydrolase (GH) families. GH2, GH29, GH86 and GH117 are the most abundant families mainly represented by β-galactosidase, α-L-fucosidase, β-agarase and α-1,3-L-neoagarooligosaccharide hydrolase. 100 GHs were predicted with signal peptide sequences indicating 61% of GHs target to the cell membrane or can be exported outside of the cell. Extracellular and membrane associated GHs may hydrolyze large extracellular polymers that cannot otherwise enter the cell. Four porins and nine sugar transporters of the major facilitator superfamily were also present in the genome which may help with the acquisition of carbohydrate molecules by the cell.

#### 3.5.2 Sulfatase analyses

It has been shown that *Kiritimatiellaeota* as well as PVC superphylum have large numbers of copies of sulfatases in their genomes (van Vliet, 2019), and it is the same case in strain NLcol2^T^. We found 165 sulfatase genes (PF00884), comprising 4.6% of all genes in the genome. Sulfatases were classified into 22 subfamilies in the SulfAtlas database, all of which belongs to family S1 fGly-sulfatases (**Table S4**). The most abundant subfamilies (>5% of total sulfatases) are S1_16, S1_7, S1_15, S1_24, S1_8, S1_19, S1_17 and S1_20. Homologous sulfatases with known enzymatic activities within these subfamilies include D-galactose-6-sulfate 6-O-sulfohydrolase, endo-/exo-xylose-2-sulfate-2-O-sulfohydrolase, endo-/exo-galactose-4-sulfate-4-O-sulfohydrolase, endo-3,6-anhydro-D-galactose-2-sulfate-2-O-sulfohydrolase, exo-fucose-2-sulfate-2-O-sulfohydrolase etc., and the known substrates of these sulfatases include alpha-/iota-/kappa-carrageenan, fucan, ulvan etc. (**Table S4**). These results imply that strain NLcol2^T^ has the potential to target a vast variety of sulfated polysaccharides, similar to isolates *K. glycovorans* (29), *P. desulfatans*, and *P. sulfatireligans* (31, 32). However, due to limited number of characterized fGly-sulfatases, there are still many unknowns about the specific substrates and/or reactions catalyzed by sulfatases in each subfamily (Barbeyron et al., 2016). 128 sulfatases have the best match genes from organisms in the PVC superphylum and 32 from *Bacteroidetes*. 96% of sulfatases (158) were predicted to have a signal peptide sequence, indicating most sulfatases could be membrane-anchored or exported outside of the cell. There are also five genes encoding formylglycine-generating enzyme and one encoding anaerobic sulfatase-maturing enzyme, which are essential for the activation of sulfatase by post-translational modification (21, 22).

#### 3.5.3 Growth on macroalgal polysaccharides

We further confirmed the ability of strain NLcol2^T^ to grow on different macroalgal polysaccharides in live cultures. Bacterial growth was observed in anaerobic cultures with agarose, agar, and iota-carrageenan, but not fucoidan. The agar used in cultivation (Difco Agar, Granulated) contains 1.7% of sulfate (Difco™ & BBL™ Manual, 2nd Edition). This is the first strain reported with the ability of utilizing agar as substrate in the *Kiritimatiellaeota* phylum. We further tested their growth on seaweeds and interestingly, cells also exhibit consistent growth on dried red algae (*Porphyra* spp.) and dried brown algae (*Saccharina japonica, Saccharina latissima*), but not on the giant kelp (*Macrocystis pyrifera*).

Genes involved in the degradation pathways of agar, iota-carrageenan and fucoidan were found in the genome (**Figure 4; Table S5**). β-agarases, ι-carrageenanases, fucoidanases and associated sulfatases were found to be key enzymes potentially degrading these macroalgal polysaccharides into monosaccharides, which then can be directed to the central metabolism for energy. 28 CAZyme gene clusters (CGC) were predicted with at least one CAZyme and one transporter present (**Table S6**). Six CGCs were also identified with a transcription factor present. 15 CGCs were identified with CAZymes as GHs, which could be putative polysaccharide utilization loci (PULs) for degrading macroalgal polysaccharides such as agar, carrageenan, and fucoidan.

**Figure 4.**
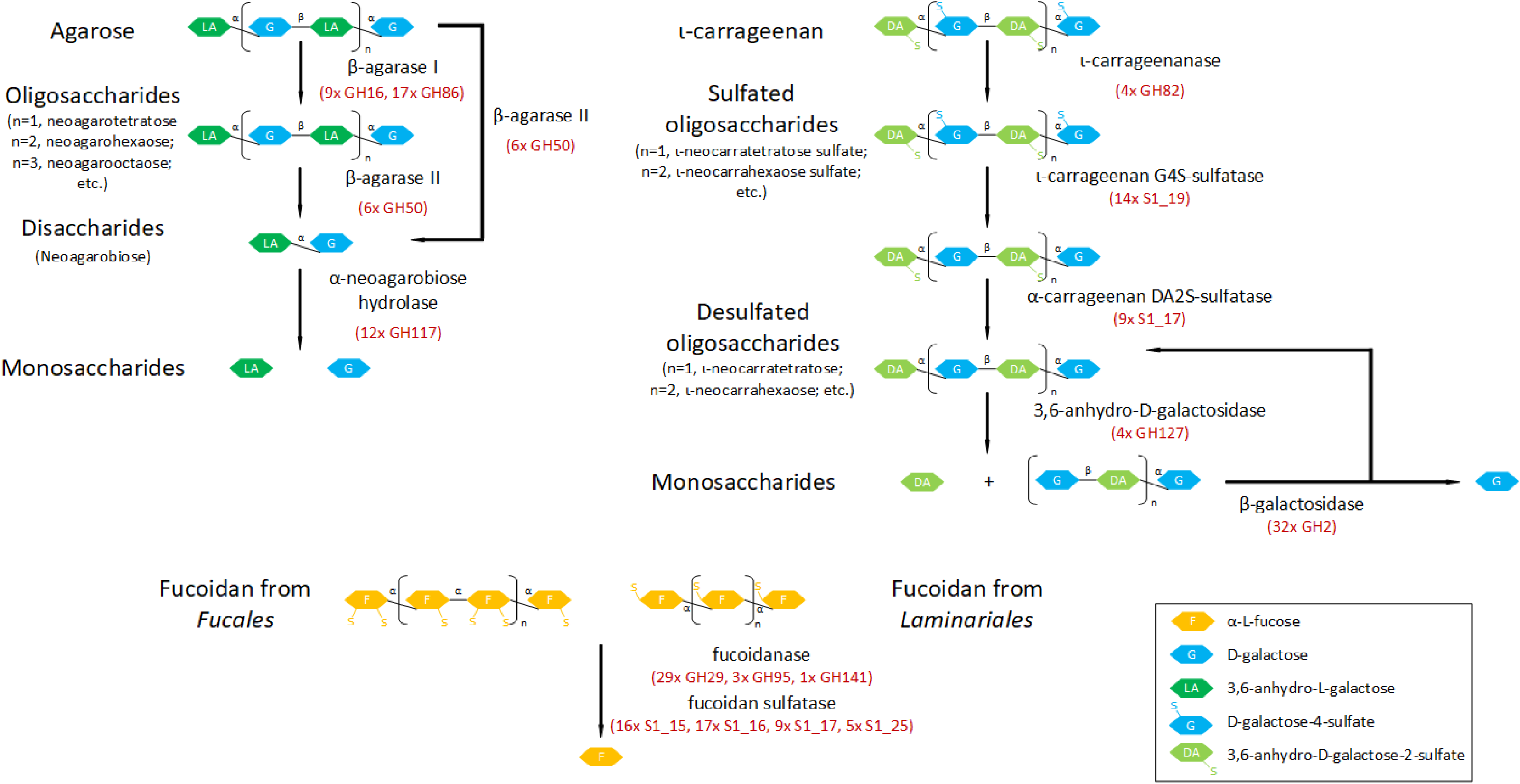
Key enzymes of found in the metabolic pathways of agar, iota-carrageenan, and fucoidan degradation. The number of gene copies are also listed. GH, glycoside hydrolase; S1_#,sulfatase classified in S1 family_subfamily.

A neighborhood analysis of the genome shows that GHs and sulfatases are often located nearby, suggesting certain sulfatases and glycoside hydrolases could be regulated together to degrade sulfated polysaccharide (16). In some cases, histidine kinase (PF07730, PF02518), response regulator (PF00072), and TonB-dependent transporters (PF00593, PF03544) are in the neighborhood too. The histidine kinase and response regulator together form a two-component signal transduction system that may help bacteria sense available substrates and respond to the changing environments (82). There are cases when sulfatases themselves cluster together, for example 4 or 6 copies in a row. A complete pathway for assimilatory sulfate reduction is also present in the genome and the cells may utilize the cleaved sulfate group for biosynthesis of reduced sulfur compounds.

A comparative study of GHs and sulfatases in selected genomes of the *Kiritimatiellales* order revealed that not all genomes harbor enzymes involved in degradation pathways of agar, iota-carrageenan, and fucoidan, and some bacteria don’t have any GHs or sulfatases at all (**Figure 1**). However, certain genomes in the *Pontiella* genus show a relatively larger component of GHs and sulfatases. This indicates that these bacteria may adopt the lifestyle of utilizing macroalgal polysaccharides like agar, carrageenan, and fucoidan as carbon and energy sources, while other clades in the *Kiritimatiellales* order may specialize on other substrates. An interesting thing to note is that some genomes in the *Pontiella* genus (on the bottom of figure 1) do not have high number of GHs or sulfatases either. This may indicate that these carbohydrate-related genes could be laterally transferred into the *Pontiella* genus but some were lost during evolution living in the environments where other available substrates were available and preferred.

### 3.6 Nitrogen fixation

We further tested nitrogen-fixing ability in live cultures of strain NLcol2^T^. The strain was able to grow on nitrogen gas as the sole N source in a nitrogen-free medium. No growth was observed when nitrogen was replaced by helium in the headspace. Bacterial growth was also supported by ammonium but not nitrate (**Figure S4**), and neither assimilatory nor dissimilatory nitrate reductase was present in the genome. Nitrogenase activity was detected by acetylene reduction assay. The production of ethylene from acetylene during bacterial growth on nitrogen gas as the sole nitrogen source showed that the cultures expressed active nitrogenases and could fix nitrogen gas into bioavailable forms to support their growth (**Figure 5**). This nitrogen-fixing ability may give them the advantage to survive in nitrogen-limiting environments.

**Figure 5.**
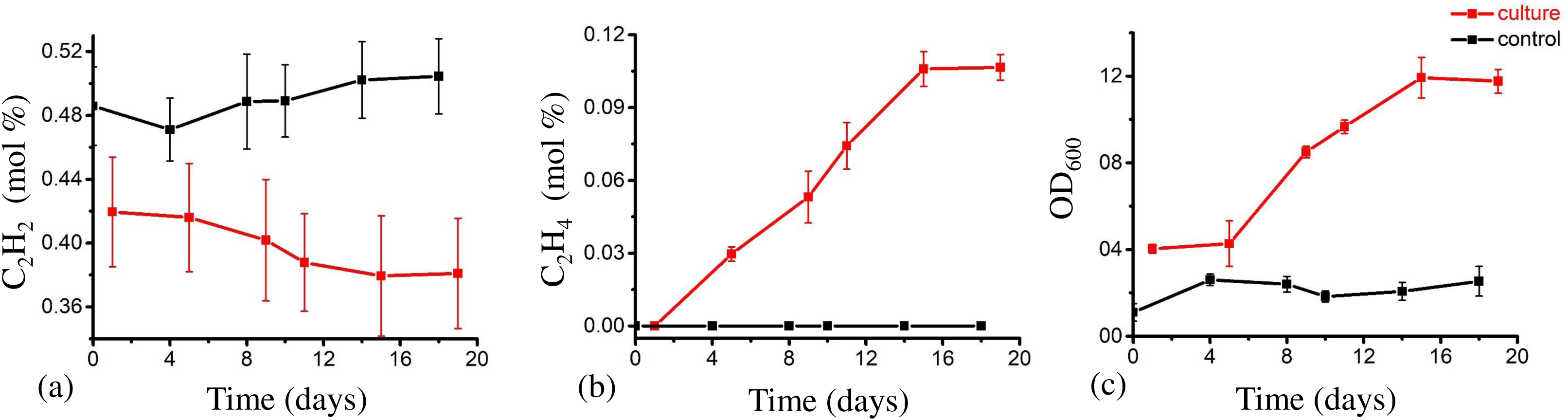
Acetylene (a), ethylene (b) percentage and OD_600_ (c) change with time in liquid culture (galactose as substrate) with nitrogen gas as the sole N source. Cultures with bacteria are labeled in red while media without inoculum (controls) are labeled in black.

Genes encoding both nitrogenase iron protein (*nifH*, PF00142) and nitrogenase molybdenum-iron protein alpha and beta subunits (*nifDK*, PF00148) are present in the genome, which together form a complete pathway of nitrogen fixation. No alternative vanadium-iron nitrogenase or iron-only nitrogenase was found. In addition to *nifHDK*, both *nifB* and *nifE* involved in the biosynthesis of nitrogenase MoFe cofactor are present in the genome. Two genes coding for nitrogen regulatory protein PII were present, which are important for the regulation of nitrogen fixation in response to nitrogen source availability (83). The rop-like protein is uncharacterized but often found in nitrogen fixation operons and may play a role in regulation (84). There are various other *nif* genes present in other parts of the genome including *nifA, M, S, U, V* which together may help regulate the function of nitrogenase (**Table S7**).

There has been no report or discussion about nitrogen fixation by bacteria in the *Kiritimatiellaeota* phylum, but we found 5 out of 53 available genomes in this phylum housing a *nifH* gene. Three were from cultivated strains F1, F21, and S94, and two were from the marine sediments at the hydrothermal vent of South Mid-Atlantic Ridge (SZUA-380 and SZUA-494). All *nifH* genes in this clade were classified as cluster III, which is dominated by distantly related obligate anaerobes (85). All 6 *nifH* genes from the *Kiritimatiellaeota* phylum form a monophyletic clade with a bootstrap value of 89 (**Figure S5**). They also cluster together with sequences from *Chlorobi*, *Bacteroidetes*, and *Delataproteobacteria* (mainly the *Desulfovibrio* genus), *Spirochaetes*, and some *Verrucomicrobia* to form a monophyletic clade with a bootstrap value of 85. This suggests that there could be lateral gene transfer between the *Kiritimatiellaeota* phylum and other phyla in this clade, but some bacteria in the *Kiritimatiellaeota* phylum may have lost *nif* genes during evolution. Nitrogen fixation genes in a methanotrophic *Verrucomicrobial* isolate *Methylacidiphilum fumariolicum* strain SolV resemble those from the *Gammaproteobacteria* which supports their acquisition of *nif* genes through lateral gene transfer (35).

## 4 Conclusion

In this study, we reported a novel anaerobic bacterial strain NLcol2^T^ isolated from microbial mats in marine sediments as the representative of a novel species in the *Pontiella* genus, which is the fifth cultivated strain in the *Kiritimatiellaeota* phylum. It represents the first strain to utilize agar as substrate with nitrogen-fixing ability in the *Kiritimatiellaeota* phylum. An extensive list of CAZymes and sulfatases shows its potential to degrade diverse macroalgae-derived sulfated polysaccharides in marine environments.

## Description of *Pontiella agarivorans* sp. nov

*Pontiella agarivorans* (a.ga.ri.vo’rans. N.L. neut. n. *agarum* agar, algal polysaccharide; L. pres. part. adj. *vorans* devouring, consuming; N.L. part. adj. *agarivorans* agar-devouring).

Cells are Gram-negative, anaerobic, non-motile cocci with a diameter of 1 μm. No spore formation was observed. Cells divide by binary fission. Colonies on agar plates are milky or ivory, circular, and smooth. Growth occurs at 10-37 °C (optimum 31 °C), with NaCl concentration between 10-60 g/L (optimum 25-30 g/L), and with pH 6.0-9.0 (optimum pH 8.0) when D-galactose was utilized as the substrate. The following substrates support growth: D-glucose, D-galactose, D-fructose, D-mannose, D-mannitol, D-xylose, D-cellobiose, lactose, sucrose, maltose, xylan, agarose, agar, iota-carrageenan, and fucoidan. The following compounds do not support growth under laboratory conditions: L-fucose, L-rhamnose, D-arabinose, meso-inositol, starch, cellulose, or alginate. The non-gaseous fermentation products from D-galactose are succinate, acetate, and malate (traces). Both ammonium and nitrogen gas can be utilized as nitrogen sources, but nitrate and nitrite were not utilized. Major cellular fatty acids are C18:0, *i-*C12:0, and *i-*C18:0.

The type strain NLcol2^T^ (= DSM 113125^T^ = MCCC 1K08672), was isolated from microbial mats on the surface of marine sediments offshore Santa Barbara, California. Genome of the type strain is 4.4 Mbp in size and DNA G+C content is 52.4 mol%. The GenBank accession number for the full-length 16S rRNA gene sequence of strain NLcol2^T^ is OQ749723, and the genome of strain NLcol2^T^ was deposited at NCBI under the accession number JARVCO000000000.

## Acknowledgments

We would like to thank Frank Kinnaman and Christoph Pierre at UC Santa Barbara for collecting sediment samples. We thank the Biological Electron Microscopy Facility at UC Davis for TEM imaging. The sequencing was carried out at the DNA Technologies and Expression Analysis Cores at the UC Davis Genome Center. We thank Professor Alex Sessions at Caltech who helped with identifying fatty acid profiles on GC-MS. We also thank the ACCESS PSC Bridges-2 for computation resources.

---

**Figure S1.**
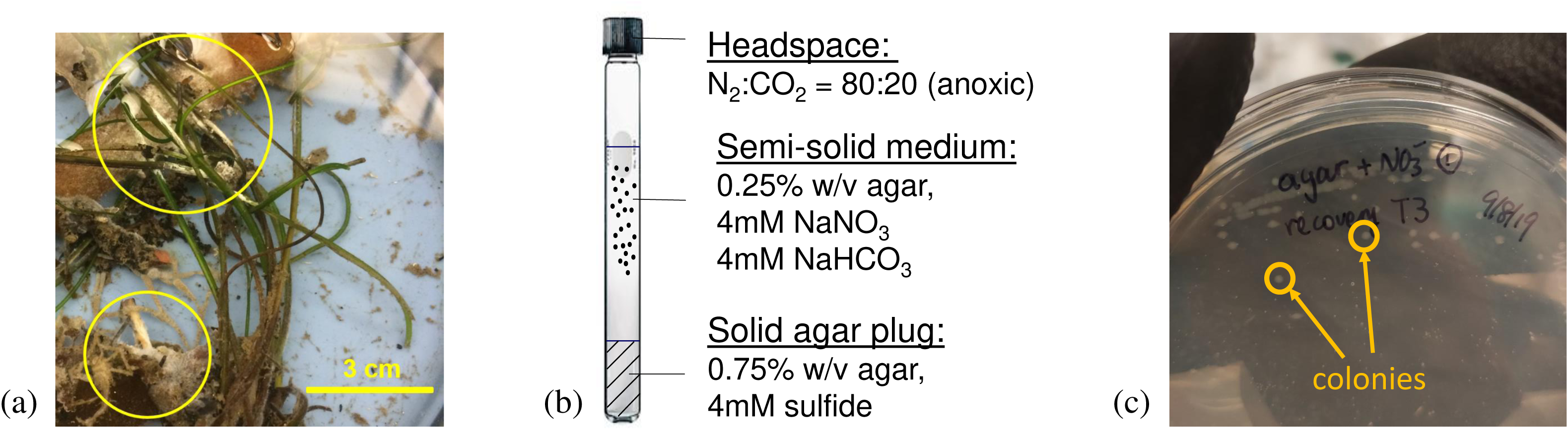
(a) Microbial mats collected on the surface of marine sediments were used as inoculum. (b) Anoxic enrichment culture setup with agar gradient seawater media. (c) Isolation of culturable strains on agar plates. Colonies were highlighted in yellow circles.

**Figure S2.**
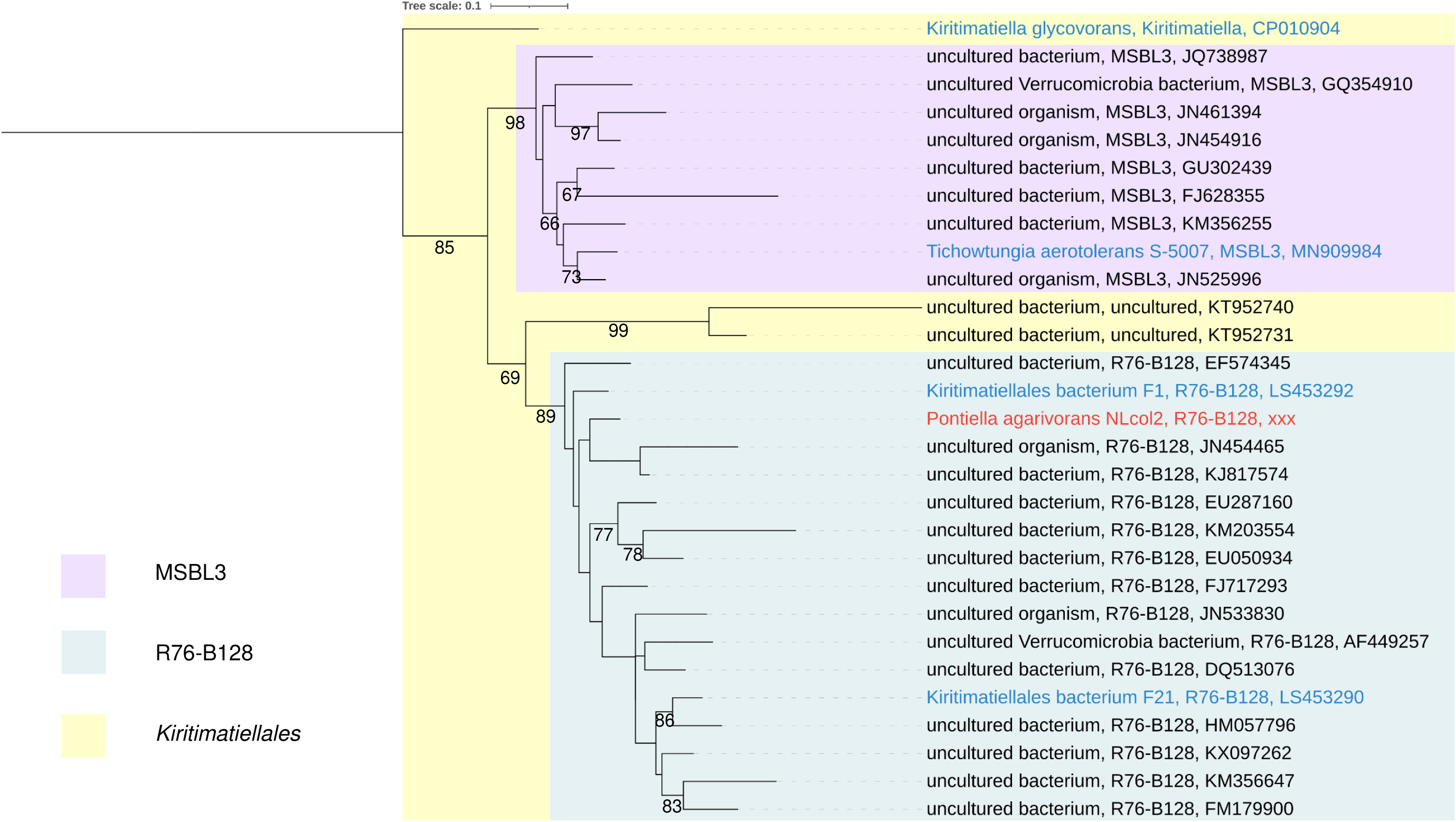
Maximum-likelihood phylogenomic tree of 16S rRNA genes of the *Kiritimatiellales*. *K. glycovorans* was also included in phylogenetic tree construction. Strain NLcol2 is labeled in red and other cultivated strains are labeled in blue. The tree was rooted with two *Verrucomicrobia* sequences (not shown) and pruned with 29 selected representative sequences. Bootstrap values over 60 are shown on the nodes.

**Figure S3.**
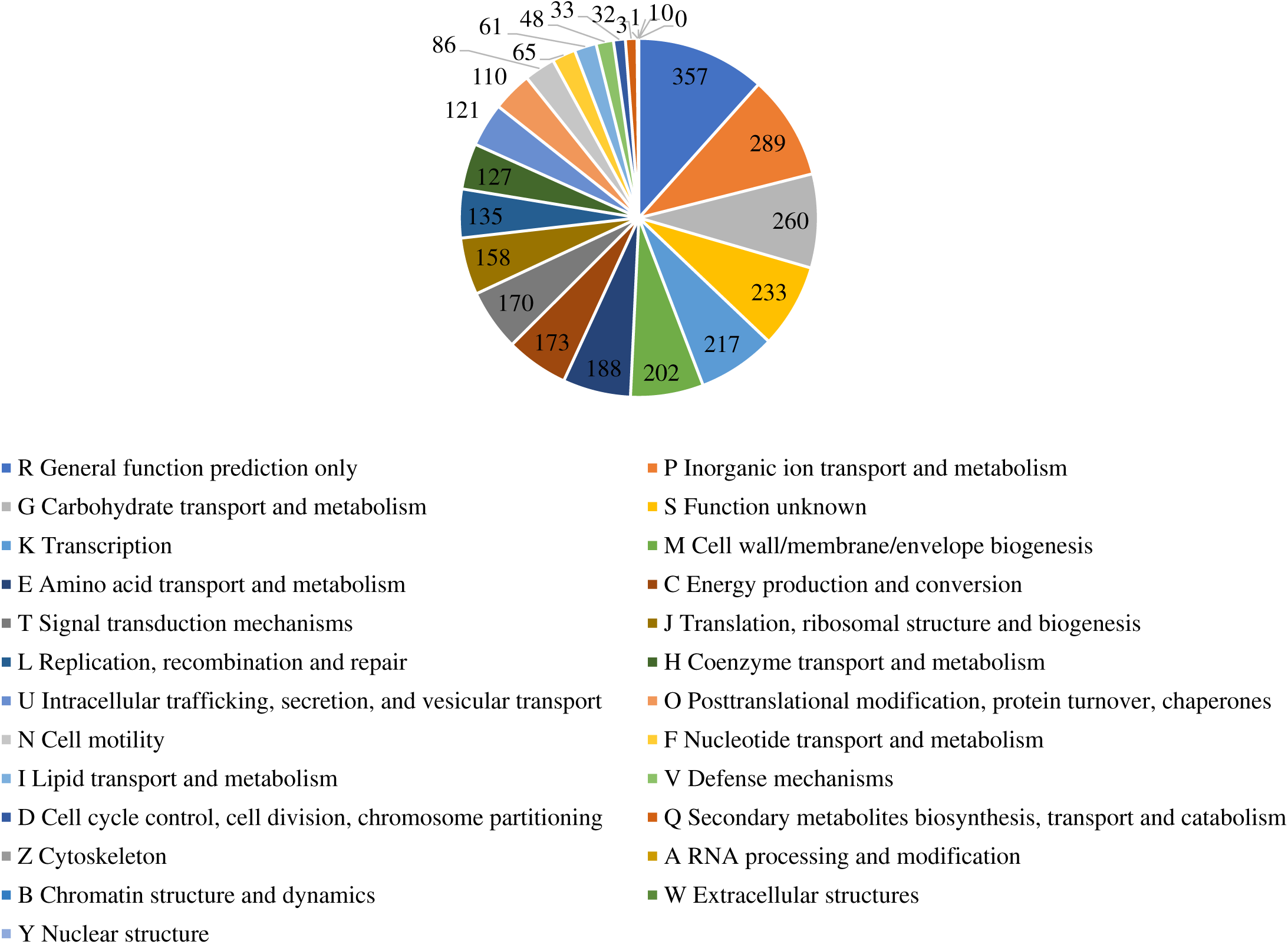
Pie chart showing number of genes assigned in COG functional categories.

**Figure S4.**
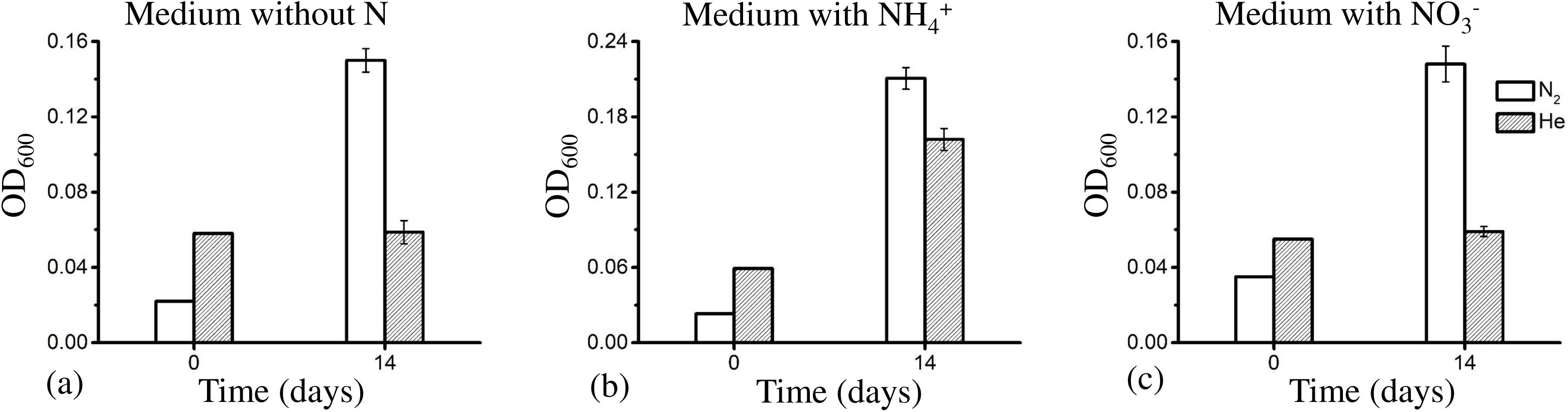
Comparisons of cultures growing in media with (a) no N, (b) ammonium or (c) nitrate supplemented. Headspace gases are nitrogen gas (white bar) or helium (grey bar).OD_600_ was measured at the beginning (day 0) and at the end (day 14).

**Figure S5.**
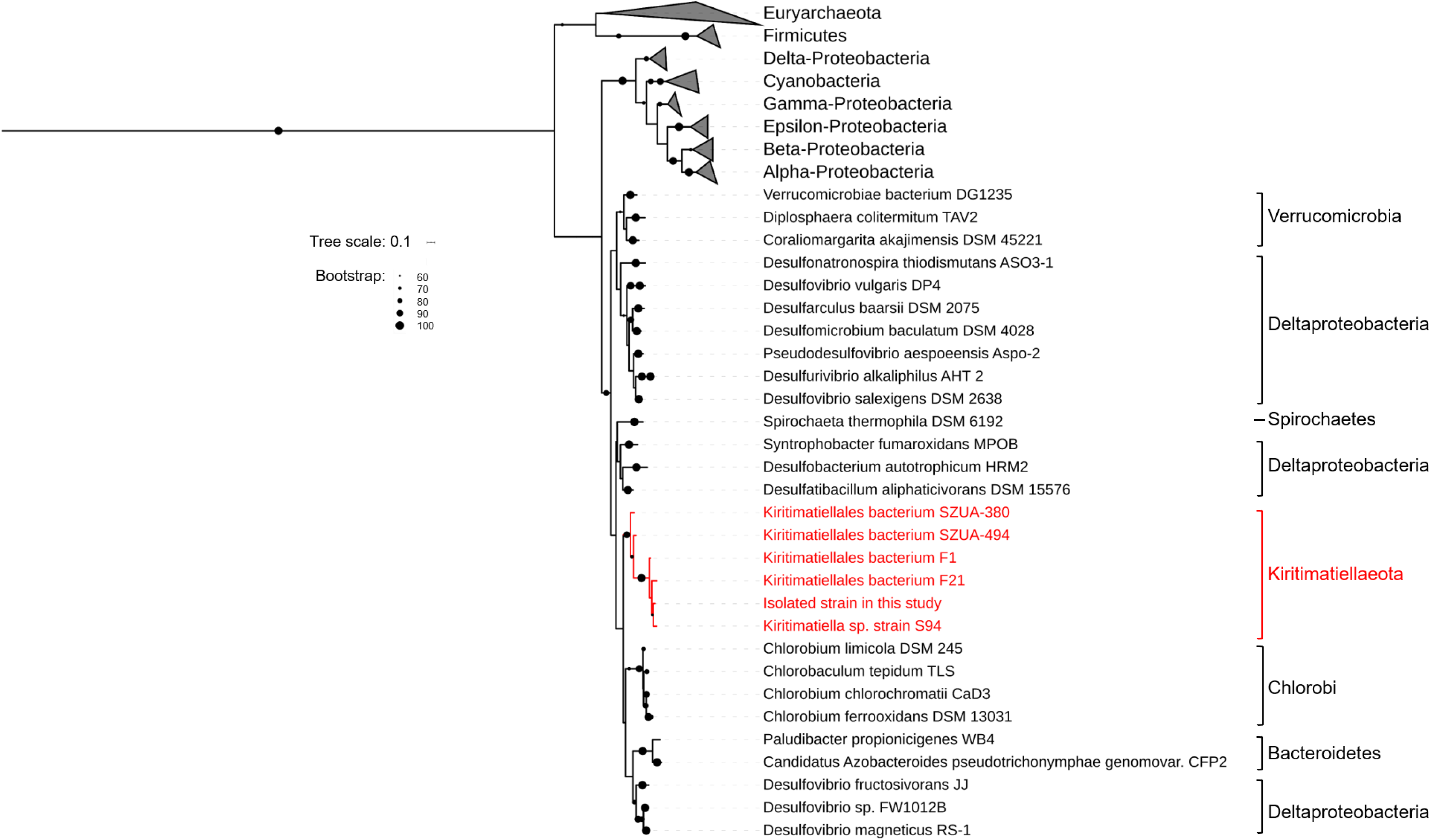
Phylogenetic placement of *nifH* genes in the *Kiritimatiellaeota* phylum on a maximum-likelihood tree. Bootstrap values over 60 are shown in dots on the nodes.

